# Nifurpirinol: A potent inducer of FSGS-like glomerular injury in transgenic zebrafish

**DOI:** 10.1101/2023.12.07.570562

**Authors:** Marianne Klawitter, Florian Siegerist, Sophie Daniel, Maximilian Schindler, Nicole Endlich

## Abstract

Identifying effective drugs for focal segmental glomerulosclerosis (FSGS) treatment holds significant importance. Our high-content drug screening on zebrafish larvae relies on nitroreductase/metronidazole (NTR/MTZ)-induced podocyte ablation to generate FSGS-like injury. A crucial factor for successful drug screenings is minimizing variability in injury induction. For this, we introduce Nifurpirinol (NFP) as a more reliable prodrug for targeted podocyte depletion. NFP showed a 2.3-fold increase in efficiency at concentrations 1600-fold lower compared to MTZ-mediated injury induction. Integration into the screening workflow validated its suitability for the high-content drug screening. The presence of crucial FSGS hallmarks, including podocyte foot process effacement, proteinuria, and activation of parietal epithelial cells, was observed. In this study, we demonstrate that NFP serves as remarkably effective prodrug for inducing an FSGS-like phenotype in zebrafish larvae and proves to be well-suited for a high-content drug screening, aiding in the identification of potential candidates for FSGS treatment.

## Introduction

Currently, chronic kidney disease (CKD) affects more than 10% of the global population, and the prevalence is increasing.^1^ Focal and segmental glomerulosclerosis (FSGS), a rare disease that is caused by podocyte injury and loss, is diagnosed by severe histologic lesions leading to glomerular damage.^2–4^ Podocytes, a postmitotic cell type, cover the outer aspect of the glomerular basement membrane (GBM) with their interdigitating foot processes and are an essential part of the filtration barrier in the kidney.^5,6^ Various factors such as hypertension, diabetes mellitus, gene mutations, or circulating factors can induce podocyte foot process (PFP) effacement, a flattening of these tiny processes.^7–11^ These morphological changes are directly linked to the loss of the size selectivity of the filtration barrier associated with proteinuria as well as edema formation, all hallmarks of kidney disease.^12,3^ Due to the limitations in healing drugs as well as therapeutic options, end-stage renal disease is often the consequence that makes dialysis or a kidney transplantation necessary.^13–15^

Screening methods have become pivotal strategies in identifying drugs that could improve FSGS.^16,17^ Our team has recently developed a high-content drug screening assay which uses transgenic zebrafish larvae as animal model. Hereby, we identified potential drugs/small molecules that showed a beneficial effect on the development of FSGS in these zebrafish larvae.^18^

The zebrafish is a well-known animal model to study renal function *in vivo.*^19–22^ The simple kidney, the pronephros, is composed of one glomerulus combined with two tubules, and shows a high similarity in morphology and protein expression compared to glomeruli of mammals.^23,24^ Moreover, the glomerular filtration starts already after 48 hours post fertilization (hpf).^24^ Since the transgenic larvae are fully transparent, fluorophores can be used as readout systems. In the zebrafish strain used for the screening, the 78 kDa eGFP-vitamin D binding protein is expressed in all vessels and mCherry in podocytes indicating a leakage of the filtration barrier as well as the loss of podocytes in living animals.^25,26^ All this taken together makes zebrafish larvae an ideal model to investigate glomerular filtration *in vivo* and surpassing the conventional cell culture systems often used for screening experiments.^16,17^

Our research group has previously established a zebrafish larval model to simulate focal segmental glomerulosclerosis (FSGS) for an *in vivo* high-content drug screening.^27^ We used a transgenic zebrafish strain called "*Cherry*" that selectively expresses the bacterial enzyme nitroreductase (NTR) and the fluorescent marker mCherry exclusively in podocytes.^26^ The NTR enzyme converts the prodrug metronidazole (MTZ) which is added to the tank water, into a cytotoxic agent that induces exclusive apoptosis of podocytes. This method is elegant, however the MTZ concentration required to induce podocyte damage is comparatively high. We also found that the extent of podocyte damage varies inter-individually, which complicates the screening process due to statistical problems when only a small number of larvae are used.

Since it was described in the literature by Bergemann *et al.* that the prodrug Nifurpirinol (NFP) is a potent and reliable prodrug for the bacterial nitroreductase working at low concentrations, we were inspired to test NFP as a potential alternative in our transgenic FSGS-like zebrafish larval model.^28^

## Materials and methods

### Zebrafish husbandry

Zebrafish were bred and maintained as previously described.^29^ The stages are given in *days post fertilization* (dpf). Three different zebrafish strains were used for this work. For observation, counting and imaging approaches, we worked with a Tg(nphs2:GAL4-VP16); Tg(UAS:Eco.nfsB-mCherry) strain, which expresses the bacterial nitroreductase and mCherry exclusively in podocytes.^26^ This strain will be referred to as *Cherry* throughout the paper. In order to demonstrate proteinuria in the screening experiments, we crossed *Cherry* larvae with a Tg(−3.5fabp10a:gc-eGFP) strain.^25^ The resulting *ScreeFi* strain additionally expresses a 78 kDa circulating vitamin D-binding fusion protein with eGFP, that is unable to pass a healthy filtration barrier. Every larva was scored for homogeneous expression of mCherry or/and eGFP before starting the treatment. A wild-type strain (ABTü) served as control. All experiments are according to the German animal protection law, were overseen and approved by the “Landesamt für Landwirtschaft, Lebensmittelsicherheit und Fischerei, Rostock” (LALLF M-V) of the federal state of Mecklenburg-Western Pomerania

### Nifurpirinol (NFP) and Metronidazole (MTZ) treatment

At 4 dpf zebrafish larvae were separated into three groups. 0.1% dimethyl sulfoxide (DMSO, Sigma Aldrich, St. Louis, MO, USA) was added to the E3 medium of the control group.^30^ The final concentrations of 80 µM MTZ (Sigma Aldrich) and 50 nM NFP (Sigma Aldrich) in 0.1% DMSO were added to the E3 medium of the injury groups. For screening experiments and *in vivo* imaging, larvae were treated for 24 hours with DMSO, MTZ or NFP. For all further experiments, the treatment period lasted 48 hours.

### *In vivo* screening

The screening was carried out as recently described.^18^ In brief, zebrafish larvae of the *ScreeFi* strain were treated with the substances at 4 dpf for 24 hours. After washout, larvae were exposed to 0.1 mg/ml tricaine anesthesia (MS-222, #E10521, Merck, Darmstadt, Germany) and positioned laterally in a 96-well plate (Greiner Bio-One, Kremsmünster, Austria) prepared with 0.7% agarose (Biozym scientific GmbH, Hessisch Oldendorf, Germany) molds. The glomerular mCherry and vascular eGFP fluorescence was imaged using the Acquifer Imaging Machine (IM; ACQUIFER Imaging, Heidelberg, Germany) focusing on the glomerular and tail region. The same larvae were imaged 24 hours later and evaluated with a custom FIJI macro.^18^ Intensity ratios were determined and statistically analysed.

### Histology

Larvae were fixed at 8 dpf in 2% paraformaldehyde (PFA) at 4°C overnight. After dehydration in an ascending ethanol series, the tissue was infiltrated and then embedded in Technovit 7100 resin (Kulzer, Hanau, Germany) as described by the manufacturer. The 4 µm plastic sections were cut with a rotational microtome (Leica Microsystems, Wetzlar, Germany) and stained with hematoxylin and eosin (HE) according to Gill. Images were acquired with an Olympus BX microscope (Tokyo, Japan).

### Immunofluorescence staining

Zebrafish larvae were fixed at 6 dpf or 8 dpf in 2% PFA for 75 minutes at room temperature and afterwards incubated in 15% sucrose diluted in 1x Dulbecco’s phosphate buffered saline (PBS, Sigma Aldrich) at 4°C overnight. On the next day, larvae were snap-frozen in liquid nitrogen in 30% sucrose mixed 1:1 with Tissue Tek O.C.T compound (Sakura Finetek Europe, BV, Netherlands). 5 µm sections were cut with a Leica CM 1950 microtome (Leica Microsystems, Heidelberg, Germany). The sections were permeabilized with 0.3% Triton X-100 for 1 minute and washed with PBS. The primary antibodies rabbit anti-pax2a (ab229318, Abcam, Cambridge, UK) 1:1000, mouse anti-atp1a1 (Developmental Studies Hybridoma Bank, University of Iowa, IA, USA) 1:150, rabbit anti-laminin (L9393, Sigma-Aldrich) 1:35 and rabbit anti-podocin 1:200 (29040, IBL, Minneapolis, MN, USA) were incubated at 4°C overnight. Subsequently, Alexa-Fluor-488 labeled anti-mouse antibody (Invitrogen, Waltham, MA, USA) and Alexa-Fluor-674 labeled anti-rabbit antibody (Invitrogen) were used for antigen detection. Nuclei were stained with Hoechst 33342 (Sigma-Aldrich) and sections were mounted with Mowiol (Carl Roth, Karlsruhe, Germany). Images were taken with the Olympus FV 3000 system using a 20x air (0.80 NA) and a 60x water (1.20 NA) objective. Fiji, an open source platform for biological image analysis, was used for image processing and morphometric measurements.^31^ The thickness of the laminin layer within the GBM was measured orthogonally as the full width at half maximum in five randomly picked capillaries of two consecutive glomerular cross-sections per larva

### Transmission electron microscopy

Zebrafish larvae were fixed at 8 dpf with 4% glutaraldehyde, 0.1% PFA and 1% sucrose in 1x PBS at 4°C overnight and then embedded in EPON 812 (Serva, Heidelberg, Germany) according to the manufacturer’s description. Semithin (500 nm) and ultrathin sections (70 nm) were cut on an Ultracut UCT microtome (Leica Microsystems). Semithin sections were stained with Richardson’s stain. Ultrathin sections were positioned on copper grids and contrasted with 5% uranyl acetate for 15 minutes and with Sato’s lead stain for 15 minutes. Transmission electron images were acquired with a LIBRA 120 electron microscope (Zeiss, Oberkochen, Germany).

### Statistics

All statistical analyses were performed with GraphPad prism V 9.4.1 (GraphPad Software, CA, USA). After testing gaussian distribution with a Kolmogorov-Smirnov test, parametric data was tested with an unpaired t test or an analysis of variance (ANOVA) whereas the nonparametric data was analysed with a Mann-Whitney U test or Kruskal-Wallis test including Dunn’s multiple comparison test. Error Bars represent ±SD and p-values lower than 0.05 were considered statistically significant.

## Results

### NFP induces edema in *Cherry* larvae

We have previously established a FSGS-like disease model by treating the transgenic zebrafish strain *Cherry* with 80 µM MTZ.^27^ The objective of this study was to investigate whether Nifurpirinol (NFP) can be used as an alternative substrate for the bacterial NTR to injure specifically podocyte. To analyze the functionality of the filtration barrier, we have taken the edema formation as a first readout. The larvae were classified into different categories dependent on the severity of the edema after the treatment (Fig. 1A). Our grading describes four categories starting from 1 (no edema) to 4 (severe edema with bend body axis) shown in Figure 1C. Titration experiments revealed that NFP induced edema formation in 3% of the larvae (at 5 dpf) at concentrations as low as 25 nM. The amount of edemas increased up to 48.5% at 8 dpf. By using 50 nM NFP, 6.1% of the larvae developed edema at 5 dpf, rising to 66.7% at 8 dpf. A further increase of the concentration of NFP up to 100 nM induced in 75.9% of the larvae edema at 5 dpf and 89.7% at 8 dpf, respectively. We used 250 nM NFP as the highest concentration and found that 90.6% of larvae developed edema at 5 dpf and 93.8% at 8 dpf, showing that we have still the dynamic range of the prodrug.

**Figure 1:**
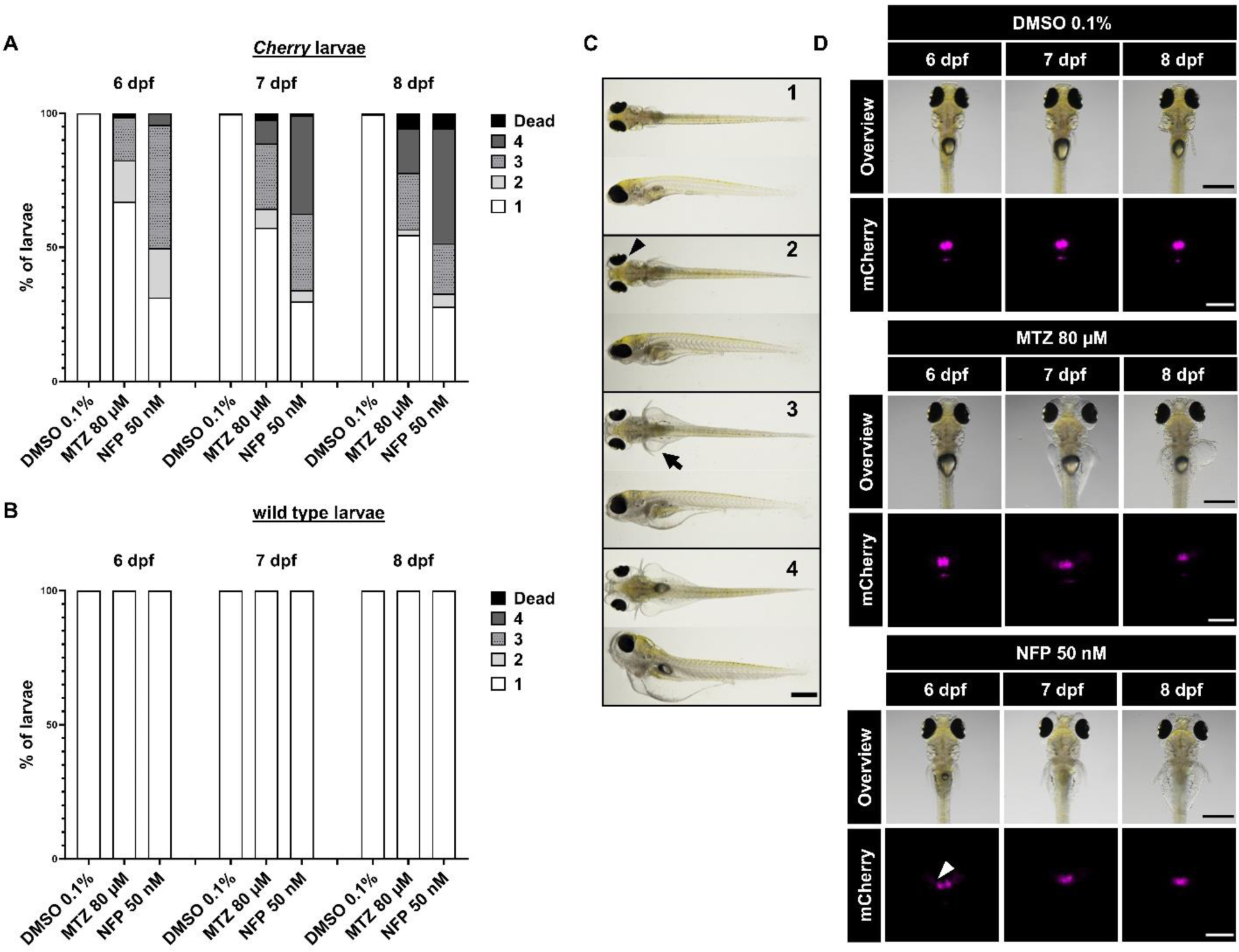
NFP induced podocyte injury and edema formation in transgenic *Cherry* zebrafish larvae. The chart represents the edema count of transgenic larvae after 48 hours of treatment at three consecutive days. At 6 dpf, 68.7% of NFP-treated and 32.9% of MTZ-treated larvae developed edema. (DMSO n=340, NFP n=326, MTZ n=337) (**A**). Wild type larvae did not respond with edema formation to NFP treatment (DMSO n=9, NFP n=25, MTZ n=24) (**B**). Exemplary images of the edematous phenotypes after podocyte depletion in dorsal and lateral orientation: (1) no edema, (2) mild edema, (3) medium edema, (4) severe edema with bent body axis, (5) dead. The arrowhead indicates slight periorbital and the arrow abdominal edema. The scale bar represents 500µm (**C**). The loss of mCherry fluorescence correlates with the severity of edema development over time. The arrowhead depicts mCherry accumulation in the proximal tubule after podocyte injury. The scale bar represents 500µm in overviews and 200µm in mCherry focused images (**D**).

### NFP is more efficient at podocyte depletion compared to MTZ

To compare the effect of NFP with the standard MTZ treatment, we administered 50 nM NFP or 80 µM MTZ to the larvae at 4 dpf as well as 0.1% DMSO to the control group (Ctrl). To assess the loss of the size selectivity of the filtration barrier and thus proteinuria, we used again the formation of edema. After 48 hours of treatment with DMSO, MTZ or NFP, the substances were washed out and the phenotype was classified for three consecutive days. At 6 dpf, 68.7% of the NFP-treated larvae and 32.9% MTZ-treated larvae developed slight to severe edema (Fig. 1A). The total number of edematous larvae per group barely increased from 6 dpf to 8 dpf, whereas the edema progressed in its severity. We found that edema formation was correlated with the reduction of the mCherry fluorescence in the glomerulus, indicating a loss of podocytes. Further, we observed an accumulation of the fluorophore mCherry in the proximal tubules due to an endocytotic uptake of mCherry (Fig. 2A). In contrast to the NFP and MTZ groups, no morphological changes were observed in the Ctrl. Additionally, we did not observe edema formation in an NTR-negative wild-type strain after the treatment with NFP or MTZ (Fig. 1B).

**Figure 2:**
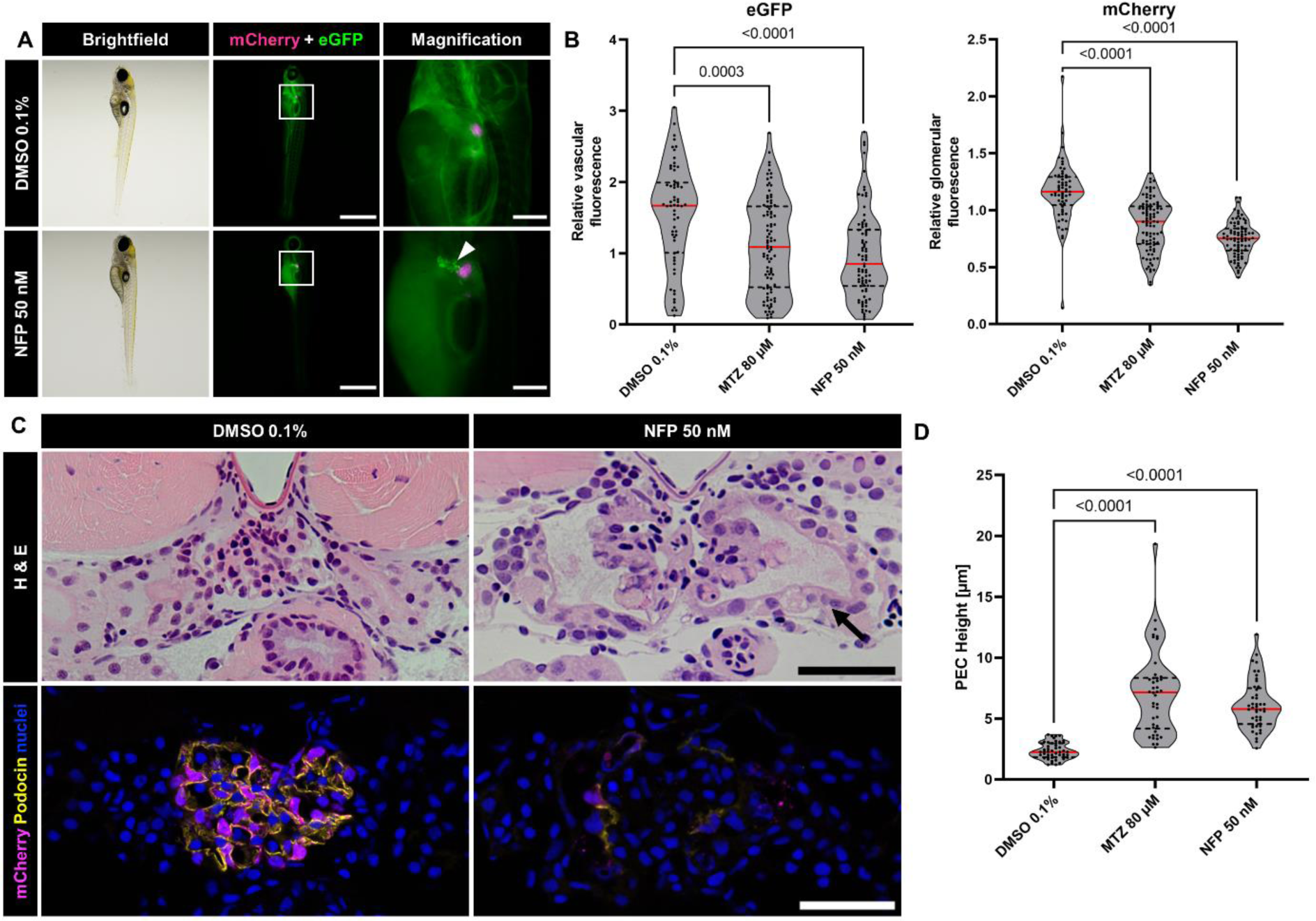
NFP induced proteinuria, podocyte depletion and histopathologic changes in the larval glomerulus. Larvae of the *ScreeFi* strain express mCherry in podocytes and a circulating 78 kDa large VDP-eGFP. The arrowhead indicates eGFP accumulation in the proximal tubule after glomerular damage with NFP. Scale bars represent 1 mm and 200 µm (**A**). The relative fluorescence intensity of the podocyte mCherry and the vascular eGFP are significantly reduced after MTZ and NFP treatment (DMSO n=62, MTZ n=93, NFP n=83) (**B**). Hematoxylin and eosin staining of control and NFP-treated larvae displays histological changes in the glomerulus. The arrow shows a thickening of the PEC-layer in podocyte depleted larvae. Immunofluorescence staining indicates a depletion of mCherry-positive podocytes accompanied by a loss of the slit membrane protein Podocin in NFP-treated larvae. Scale bars represent 50 µm (**C**). Measurements of the PEC-layer demonstrate a significant increase in cellular height after podocyte depletion (DMSO n=16, MTZ n=14, NFP n=15) (**D**).

### NFP leads to a FSGS-like glomerular injury

To proof whether the treatment of the larvae with NFP instead of MTZ induces a FSGS-like phenotype, we analyzed the glomeruli after the treatment. We found that the fluorescence intensity of mCherry correlated with the expression of podocin (Fig. 2C). Moreover, we identified an activation of parietal epithelial cells (PECs) in different ways. Thus, H&E staining showed a significant increase of the height of PECs (Fig. 2C and 2D). After 48 hours of NFP treatment, mean cellular height was 2.35 µm in Ctrl (n=16) and increased to 7.11 µm in the MTZ-treated (p< 0.0001; n=14) and to 6.12 µm (p< 0.0001; n=15) in the NFP-treated larvae. Additionally, we found by immunofluorescence staining that the activated PECs were strongly Pax2a-positive (Fig. 3A). These cuboidal Pax2a*-*positive cells were found on the capillary tuft after NFP-induced podocyte depletion. Quantification of Pax2-positive cells showed an approximately 10-fold higher number of cells per µm^2^ in the MTZ group (3.2 × 10^−3^ per µm^2^ (p< 0.0001; n=24)) as well as in NFP group (2.3 × 10^−3^ per µm^2^ (p< 0.0001; n=25)) in contrast to the Ctrl group (3.2 × 10^−4^ per µm^2^ (n=25)).

**Figure 3:**
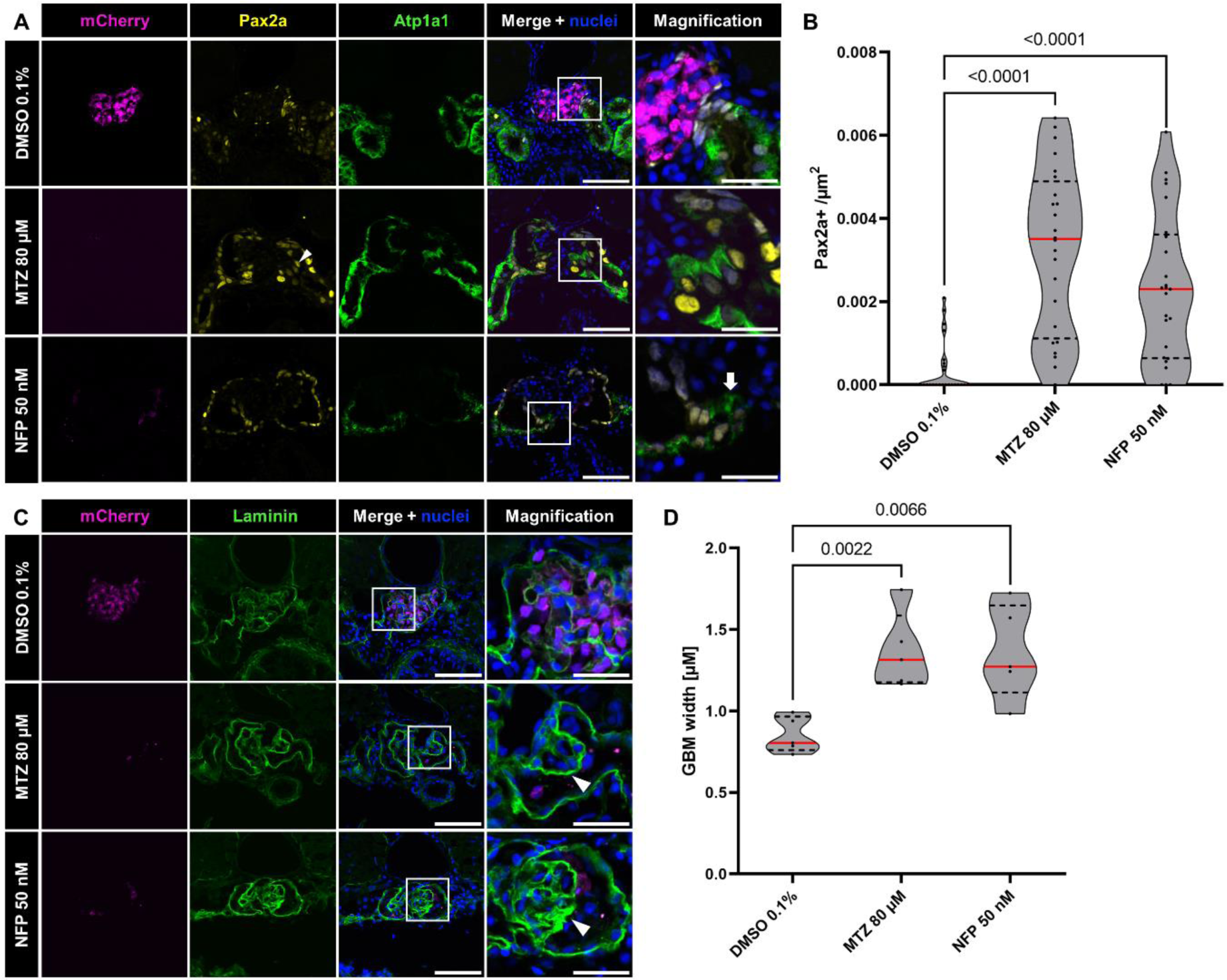
Immunofluorescence staining reveals various glomerular changes in podocyte depleted larvae. Representative confocal micrographs of MTZ and NFP-treated larvae show Pax2a (arrowhead) and Atp1a1-positive (arrow) cells on the PEC-layer and the glomerular tuft. Scale bars represent 50 µm and 20 µm (**A**). Podocyte depletion results in an increase of Pax2a-positive cells on the glomerular tuft (DMSO n=25, MTZ n=24 and NFP n=25) (**B**). Immunofluorescence staining for laminin shows partial thickening of the GBM (arrowheads) in NFP and MTZ-treated groups. Scale bars represent 50 µm and 20 µm (**C**). The thickness of the Laminin layer within the GBM was significantly increased after partial podocyte depletion. (DMSO n=10, MTZ n=10, NFP n=10) (**D**).

Another indicator of activated PECs is the staining for the Na+K+-ATPase subunit alpha 1 (Atp1a1). We found a strong expression of Atp1a1 on the parietal leaf of the Bowman’s capsule and on the glomerular tuft in contrast to the Ctrl group where no Atp1a1 signal was visible in the glomeruli (n=25 per group (Fig 3A)).

Since matrix accumulation is a further hint for FSGS, we stained the histological sections with an antibody against laminin and determined the thickness of the GBM. Figure 3C and D show that the width of the GBM is 1.37 µm (p= 0.0022; n=5) in the MTZ group and 1.36 µm (p= 0.0066; n=5) in NFP group in contrast 0.85 µm (n=5) in the Ctrl. Ultrastructural analysis by using transmission electron microscopy (TEM) showed a severe PFP effacement in the NFP-treated larvae as well as in the MTZ-treated larvae (Figure 4 A’, A’’). Additionally, we found podocyte with pseudocysts and microvillous formation and the formation of tight junctions between different podocytes, as shown in Figure 4 A’ and B’’. We also found PECs that have had cilia as well as a basal accumulation of mitochondria (Fig 4B’, B’’).

**Figure 4:**
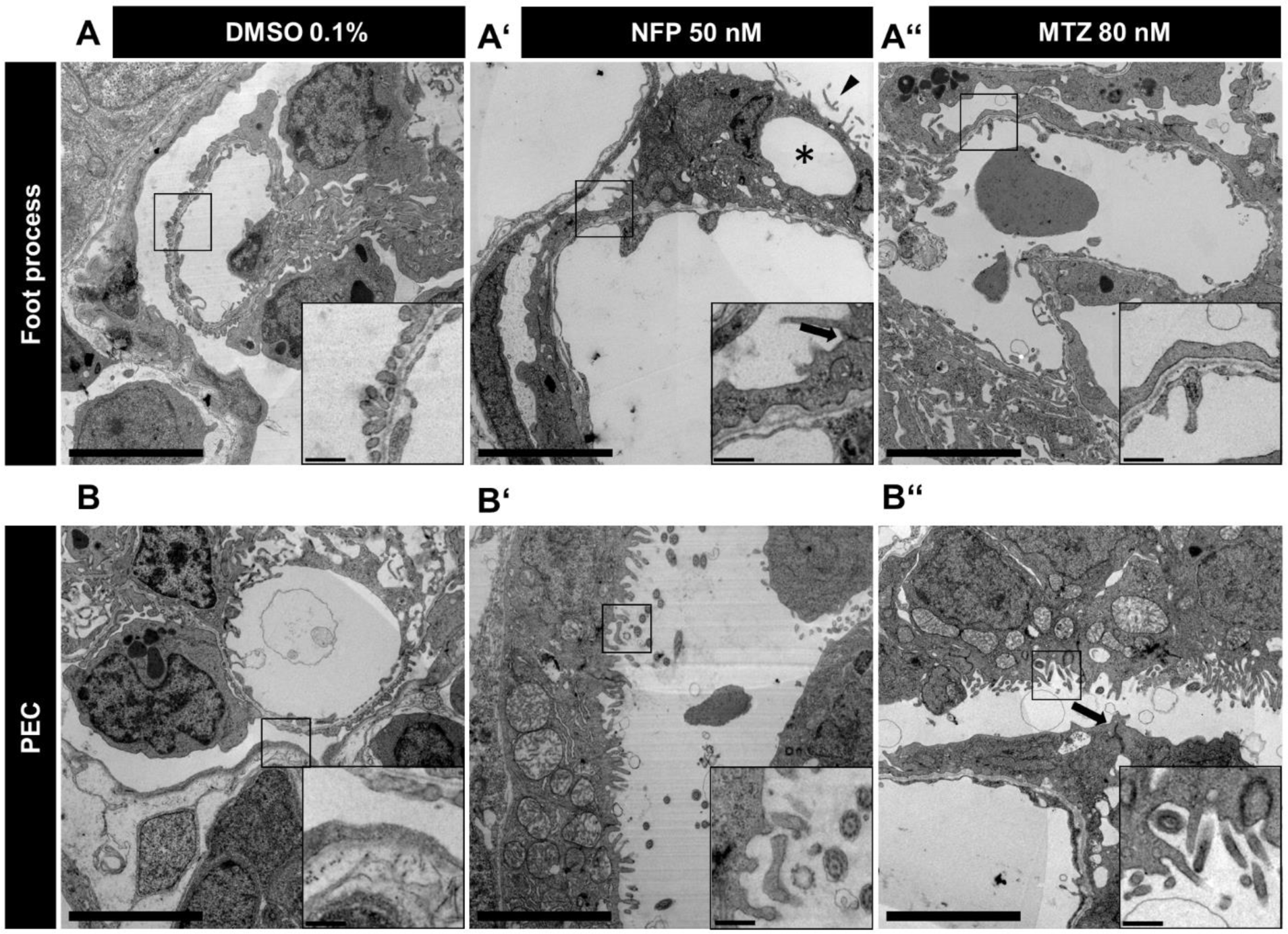
Transmission electron micrographs show pronounced foot process effacement and an activation of PECs. Transmission electron micrographs of DMSO control larvae represent a healthy glomerular capillary. The filtration barrier consists of a fenestrated endothelium and a three layered GBM on which interdigitating podocyte foot processes are connected to form the slit diaphragm (**A**). Podocytes develop intracellular pseudocysts (asterisk), apical microvillous formations (arrowhead) and tight junctions (arrow) after NFP treatment. The magnification points out severe foot process effacement (**A’**). A podocyte depleted glomerular capillary after MTZ treatment shows similar severe foot process effacement (**A’’**). The PEC-layer of the Bowman’s capsule in healthy controls shows a flat monolayer epithelium (**B**). Activated PECs display an increased cellular height after NFP treatment. Apical development of cilia and basal accumulation of mitochondria is frequently visible in the PECs of all podocyte depleted larvae (**B’**). A similar activation of PECs is visible in MTZ-treated larvae. The arrowhead accentuates tight junctions between podocytes. Scale bars represents 5 µm in overviews and 0.5 µm in magnifications (**B’’**).

Taken together, we identified activated PECs in partial podocyte-depleted larvae, characterized by increased height and strong Pax2a positivity, as well as notable changes in Atp1a1 expression and increased GBM thickness, suggesting matrix accumulation.

### NFP is well suited for high-content drug screening

To find out whether NFP instead of MTZ could be used in the high-content screening, the fluorescence intensities of the two used readouts of the screen were determined. As shown in Figure 2B, the fluorescence intensities of the vascular eGFP and the mCherry signal of the podocytes were automatically measured in a 96-well plate under screening conditions on two consecutive days. The ratio was determined as the fluorescence intensity changes between day 5 and day 6. The vascular fluorescence ratio in the Ctrl was 1.54 (SD=0.72; n=62) and decreased to 1.11 due to the MTZ treatment (80 µM, p= 0.0003; SD=0.65; n=93). By using 50 nM NFP, we found a reduction of the eGFP ratio to 0.96 (p< 0.0001; SD=0.61; n=83). The mCherry ratio of the untreated larvae with 1.16 (SD=0.27) decreased to 0.87 (p< 0.0001; SD=0.23) in the MTZ-treated group. In the NFP-treated larvae, the ratio was 0.75 (p<0.001; SD=0.15) indicating a stronger loss of the mCherry fluorescence when compared to the MTZ-treated group by using a significant lower concentration.

### Specific histone deacetylase (HDAC) inhibitors prevented edema formation in the NFP-induced FSGS-like model

Since pan-HDAC inhibitors such as Belinostat, Oxamflatin and Panobinostat were shown to impair edema formation in MTZ-treated larvae, we wanted to analyze whether these inhibitors also show beneficial effects on NFP-treated zebrafish larvae.^18^

After the co-treatment of NFP (50 nM) with Belinostat (100 µM), the zebrafish larvae were classified into edema categories. All abnormalities of the larval zebrafish with clear absence of periorbital edema were considered as malformations (Fig 5D). The administration of NFP induced edema in 58.2% of the larvae at 6 dpf, in contrast to the NFP and Belinostat co-treatment group, which exhibited only mild edema in 1.8% of the larvae at 6 dpf. 24 hours after the washout of both substances, the rate of mild to moderate edema increased to 50.1%.

**Figure 5:**
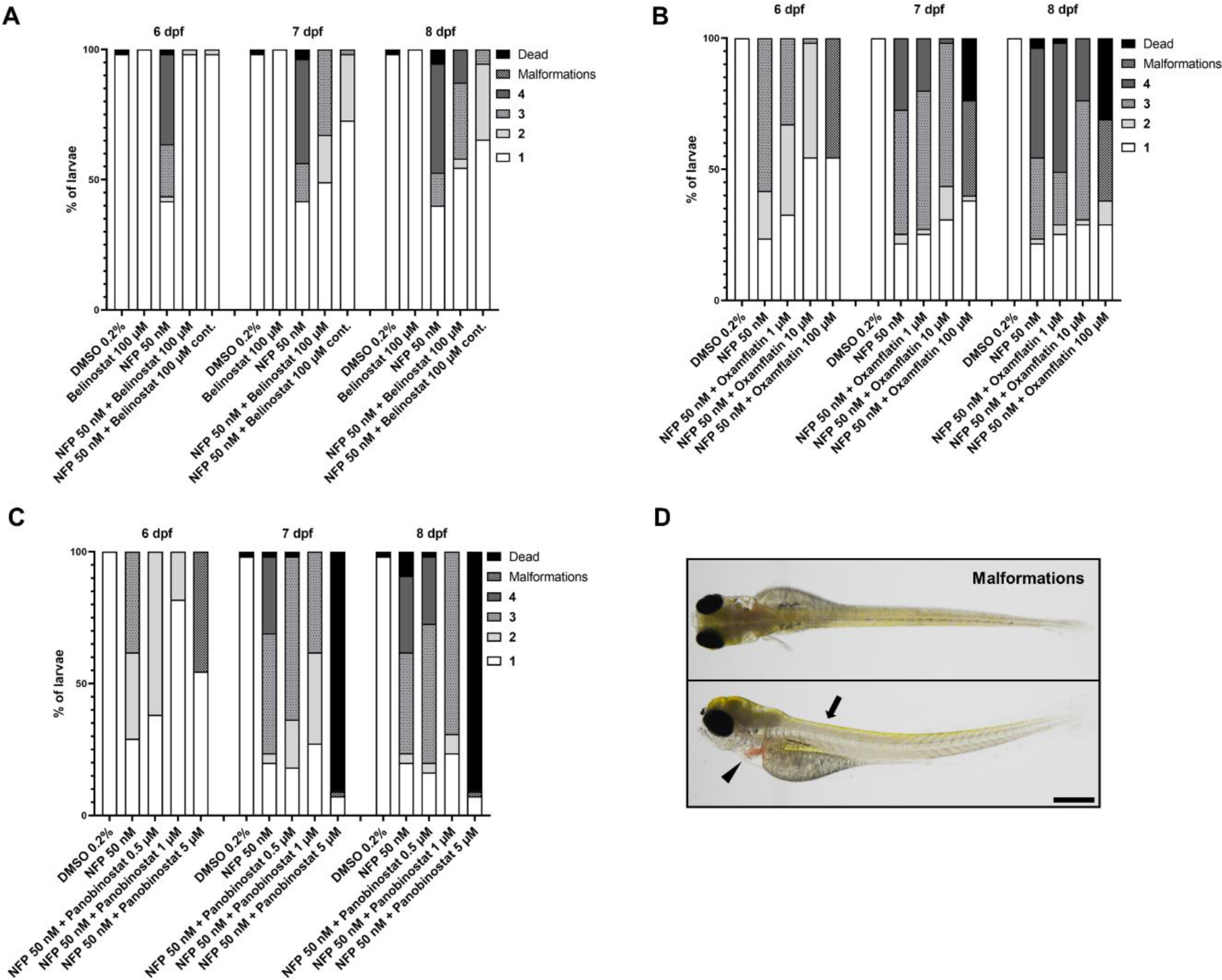
Treatment with HDAC inhibitors showed a reduction in edema progression. *Cherry* larvae were treated with Belinostat and NFP for 48 hours and edema severity was classified for three consecutive days. The simultaneous treatment showed a substantial reduction in formation and edema severity. 100 µM Belinostat has had no toxic effect alone. Continuous treatment with 100 µM Belinostat until 8 dpf enhanced the beneficial effect compared to the washout group (**A**). Treatment with Oxamflatin showed a clear protective effect at concentrations up to 10 µM. Treatment with 100 µM led to pronounced malformations. (**B**). Treatment with Panobinostat resulted in reduction of edema at concentrations up to 1 µM. A concentration of 5 µM led to formation of malformations and an increased mortality (**C**). Images of the 4 dpf zebrafish larvae showed exemplary malformations after substance treatment with a curved body axis (arrow), pericardial edema (arrowhead) with the absence of periorbital or abdominal edema. The scale bar represents 200 µm (**D**).

To find out whether a continuous treatment with Belinostat could improve the observed protective effect, we treated the larvae with 100 µM Belinostat daily, starting from 6 to 8 dpf. At 7 dpf, the percentage of zebrafish larvae with mild to moderate edema was 27.3%, increasing to 34.5% at 8 dpf. Additionally, the severity of the edema formation was significantly reduced. In contrast, the group without Belinostat showed edema in 58.1% of the larvae at 7 dpf and 60% at 8 dpf (Figure 5A).

Oxamflatin (1 µM), a structural similar compound to Belinostat, also showed a reduction of the edema formation by 9% compared to the NFP group at 6 dpf. This effect could be enhanced by using higher concentrations of Oxamflatin (10 µM, 100 µM). As shown in Figure 5B, edema formation was reduced by 31% and 45.4%, respectively to the NFP group. A third pan-HDAC inhibitor, Panobinostat, reduced the edema rate by 9% at a concentration of 0.5 µM and by 51.9% at 1 µM compared to the NFP group. Higher concentrations were not possible to use since it induces severe malformations of the larvae or led to the death of the larvae (Figure 5 C). These results clearly show the same positive effect of the pan-HDAC inhibitors Belinostat, Panobinostat and Oxamflatin in the NFP-induced FSGS zebrafish model.

## Discussion

FSGS is a rare and often life-threatening disease resulting from the damage as well as the loss of a highly specialized and postmitotic cell type, the podocytes.^3^ Current therapies involve corticosteroids and immunosuppressant therapies, with renal replacement therapies being the last option.^13^ This situation highlights the need for new, protective drugs. To address this, we developed a high-content drug screening in which transgenic zebrafish larvae were treated with Metronidazole (MTZ), a well-established prodrug that is converted into a toxin by the bacterial nitroreductase (NTR) selectively expressed in the podocytes in a transgenic zebrafish strain.^18^ After the damage and loss of podocytes, the zebrafish larvae develop a FSGS-like injury.^27^ Since MTZ, which is used at micromolar concentrations showed a high inter-individual variability in generation of FSGS, we wanted to test an alternative substrate for the bacterial nitroreductase. Bergemann and coworkers reported about an alternative prodrug for the NTR-based zebrafish model. They found that NFP, which is also a nitroaromatic antibiotic compound, had a more efficient and robust cell ablation effect in the transgenic zebrafish larvae.^28^ Comparable to the work of Bergemann et al., NFP could be used in an approx. 2000-fold lower concentration for targeted cell depletion in contrast to MTZ. In addition to the work in which experiments were performed on dopaminergic neurons, pancreatic beta cells and osteoblasts, we were able to show that efficient podocyte damage can already be achieved in the nanomolar range. In our titration experiments we found that a concentration of 50 nM NFP showed a 2.3-fold higher effect in edema formation, a hallmark of kidney damage, compared to 80 µM MTZ. Integration of the NFP damage model into the workflow of the high content screening resulted in a more valid outcome with lower standard deviations than the previous treatment with MTZ.

FSGS is characterized by different hallmarks such as edema formation, podocyte loss, matrix accumulations as well as activation of PECs.^32–34^ To proof that the morphological damage and changes after the NFP treatment are comparable to the injury observed by the use of MTZ, we studied the glomeruli and the ultrastructural changes of the treated larvae by H&E, immunofluorescence and TEM and compared them to the MTZ-induced FSGS-like injury.

In healthy kidney, PECs are found as flat cells along the Bowman’s capsule in the glomerulus and may have properties similar to those of renal progenitor cells^35,36^. Following induction of renal damage by NFP, there was a marked enlargement of these PECs, consistent with what Kuppe et al. observed in mammalian models of FSGS.^37^ In addition, these cells also expressed markers typical of proximal tubule cells together with those of PECs. This dual feature was similarly reported in human FSGS cases by Dijkman and coworkers.^38^

Since we have identified several pan-HDAC inhibitors in our recent high-content screening, we wanted to compare these results with the results received with NFP. Our experiments showed the same effects of the pan-HDAC inhibitors such as Belinostat, Oxamflatin as well as Panobinostat on the reduction of the NFP-induced edema formation in zebrafish larvae as we have seen with MTZ treatment.

Taken together, NFP is a prodrug for the bacterial reductase which can be used at nanomolar concentrations to induce a FSGS-like injury in a transgenic zebrafish strain in a more reliable way and can therefore be also implemented into the high-content screening pipeline for drugs to treat FSGS.

### Limitations of this study

In addition to its numerous advantages, the zebrafish larva as an experimental animal model in nephrology also exhibits minor limitations. Although structurally highly comparable to mammalian nephrons, it is noteworthy that the zebrafish pronephros lacks certain parts like a loop of henle and the macula densa. Despite many similarities in the genome and structure, validation experiments in rodents are essential as differences in drug metabolism need to be considered. Intravenous injections of drugs/small molecules into the larval vasculature require experience and are time-consuming. However, this problem can mostly be solved by addition of the substance to the tank water. As the zebrafish larvae are still in a developing stage, regenerative processes cannot be excluded. The various advantages of zebrafish larvae clearly outweigh minor limitations which make this model an excellent bridge between cell culture experiments and mammalian models.

## Acknowledgments

This work was supported by two scholarships of the Gerhard Domagk program of the University Medicine Greifswald to MK and SD and by a grant of the Federal Ministry of Education and Research (BMBF, grant 01GM1518B, STOP-FSGS) to NE. Additionally, the Bundesministerium für Wirtschaft und Klimaschutz (BMWi) funded this project (Grant No. 16KN077229, title: AlternaTier-vivoPod). This work was generously supported by Südmeyer fund for kidney and vascular research (“Südmeyer Stiftung für Nieren- und Gefäßforschung”) and the Dr. Gerhard Büchtemann fund, Hamburg, Germany. The authors thank Sophia-Marie Bach and Mandy Weise for technical assistance as well as Oliver Zabel and Steffen Prellwitz for excellent zebrafish husbandry.

